# The genome of the fungal pathogen *Verticillium dahliae* reveals extensive bacterial to fungal gene transfer

**DOI:** 10.1101/485037

**Authors:** Xiaoqian Shi-Kunne, Mathijs van Kooten, Jasper R.L. Depotter, Bart P.H.J. Thomma, Michael F. Seidl

**Affiliations:** Laboratory of Phytopathology, Wageningen University, Droevendaalsesteeg 1, 6708 PB Wageningen, The Netherlands; Department of Crops and Agronomy, National Institute of Agricultural Botany, Huntingdon Road, CB3 0LE Cambridge, United Kingdom

**Author notes:** These authors contributed equally to this work (Mathijs van Kooten, Jasper R.L. Depotter). These authors contributed equally to this work (Bart P.H.J. Thomma, Michael F. Seidl). Authors for correspondence: Bart P.H.J. Thomma, Laboratory of Phytopathology, Wageningen University, Droevendaalsesteeg 1, 6708 PB Wageningen, The Netherlands.; Michael F. Seidl, Laboratory of Phytopathology, Wageningen University, Droevendaalsesteeg 1, 6708 PB Wageningen, The Netherlands.

## Abstract

Horizontal gene transfer (HGT) involves the transmission of genetic material between distinct evolutionary lineages and can be an important source of biological innovation. Reports of inter-kingdom HGT to eukaryotic microbial pathogens have accumulated over recent years. *Verticillium dahliae* is a notorious plant pathogen that causes vascular wilt disease on hundreds of plant species, resulting in high economic losses every year. Previously, the effector gene *Ave1* and a glucosyltransferase-encoding gene were identified as virulence factor-encoding genes that were proposed to be horizontally acquired from a plant and a bacterial donor, respectively. However, to what extent HGT contributed to the overall genome composition of *V. dahliae* remained elusive. Here, we systematically searched for evidence of inter-kingdom HGT events in the genome of *V. dahliae* and provide evidence for extensive horizontal gene acquisition from bacterial origin.

## INTRODUCTION

Genetic information is generally vertically transferred from parents to their offspring. However, genetic information can also be transmitted laterally between reproductively isolated species, often referred to as horizontal gene transfer (HGT). It has been well established that HGT plays a significant role in the adaptive evolution of prokaryotic species (Eisen 2000; Koonin et al. 2001; Bapteste et al. 2009). Three well-characterized mechanisms contribute to DNA uptake by prokaryotes, namely transformation, conjugation and transduction. With transformation, a DNA fragment from a dead, degraded bacterium or another donor enters a competent recipient bacterium (Johnston et al. 2014), while conjugation is the active DNA transfer between prokaryotic cells by direct cell-to-cell contact or by a bridge-like connection between two cells (Norman et al. 2009). Finally, transduction involves the transfer of a DNA fragment from one prokaryotic cell to another by a virus or viral vector. HGT in prokaryotes takes place at all taxonomic levels: from individuals of the same population up to inter-kingdom transfers.

HGT also contributes to the evolution of eukaryotes, although it occurs much less frequent than in prokaryotes (Kurland et al. 2003; Bock 2010). Horizontal gene transfer has played an important role in the evolution of pathogenic traits in plant pathogens (Soanes & Richards 2014). For instance, the wheat pathogenic fungi *Pyrenophora tritici-repentis* and *Phaeosphaeria nodorum* both contain the near-identical effector gene *ToxA* that encodes a host-specific toxin that acts as pathogenicity factor (Friesen et al. 2006). Compelling evidence revealed that *ToxA* was acquired by *P. tritici-repentis* from *P. nodorum* via HGT, giving rise to pathogenicity of the former species on wheat plants (Friesen et al. 2006). Interestingly, *ToxA* was similarly found in the genome of yet another wheat pathogen, *Bipolaris sorokiniana* (McDonald et al. 2018).

Although knowledge on mechanisms of HGT in eukaryotes remains limited, HGT in filamentous plant pathogens is thought to occur more often between closely related species since donor and recipient species share similar genomic architectures (Mehrabi et al. 2011). For the filamentous plant pathogenic fungus *Fusarium oxysporum* it was demonstrated that an entire chromosome can be transferred from one strain to another through co-incubation under laboratory conditions *in vitro* (Ma et al. 2010; van Dam et al. 2017).

Intriguingly, inter-kingdom HGT also contributed to the genome evolution of filamentous plant pathogens despite the evolutionary distant relation with the donor. For example, phylogenetic analyses of plant pathogenic oomycete species revealed 34 gene families that have undergone HGT between fungi and oomycetes (Richards et al. 2011). The repertoire of HGT candidates includes genes encoding proteins with the capacity to break down plant cell walls and effector proteins for interacting with host plants (Richards et al. 2011). Furthermore, genomic and phylogenetic analyses of the fungus *Fusarium pseudograminearum*, the major cause of crown and root rot of barley and wheat in Australia, has revealed that a novel virulence gene was horizontally acquired from a bacterial species (Gardiner et al. 2012).

The fungal genus *Verticillium* contains nine haploid species plus the allodiploid *V. longisporum* (Inderbitzin et al. 2011; Depotter et al. 2017). These ten species are phylogenetically subdivided into two clades; Flavexudans and Flavnonexudans (Inderbitzin et al. 2011; Shi-Kunne et al. 2018). The Flavnonexudans clade comprises *V. nubilum, V. alfalfae, V. nonalfalfae, V. dahliae*, and *V. longisporum*, while the Flavexudans clade comprises *V. albo-atrum, V. isaacii, V. tricorpus, V. klebahnii* and *V. zaregamsianum* (Inderbitzin et al. 2011). Among these *Verticillium* spp., *V. dahliae* is the most notorious plant pathogen that is able to cause disease in hundreds of plant species (Inderbitzin & Subbarao 2014; Fradin & Thomma 2006). Furthermore, also *V. albo-atrum, V. alfalfae, V. nonalfalfae* and *V. longisporum* are pathogenic, albeit with narrower host ranges (Inderbitzin & Subbarao 2014). Although the remaining species *V. tricorpus, V. zaregamsianum, V. nubilum, V. isaacii* and *V. klebahnii* have incidentally been reported as plant pathogens too, they are mostly considered saprophytes that thrive on dead organic material and their incidental infections should be seen as opportunistic (Inderbitzin et al. 2011; Gurung et al. 2015; Ebihara et al. 2003). *Verticillium* spp. are considered to be strictly asexual, yet various mechanisms contributing to the genomic diversity of *V. dahliae* have been reported (Seidl & Thomma 2017, 2014; Faino et al. 2016), including HGT (de Jonge et al. 2012; Klosterman et al. 2011). Previously, *Ave1* and a glucosyltransferase encoding gene were found to be acquired from a plant and a bacterial donor, respectively, both of which were found to contribute to *V. dahliae* virulence during plant infection (de Jonge et al. 2012; Klosterman et al. 2011). These two inter-kingdom HGT events inspired us to study the extent and potential impact of inter-kingdom HGT to *Verticillium dahliae*.

## RESULTS

### Extensive inter-kingdom HGT to *V. dahliae* and other Ascomycete fungi

To systematically search for genes in the *V. dahliae* genome that are derived from inter-kingdom HGT, we downloaded a recent UniProtKB proteome database that contains proteomes of 5,074 species, including 4,123 bacteria, 182 archaea and 769 eukaryotes (Figure 1). The database contains proteomes of 373 fungal species, including *V. dahliae* (strain VdLs17), *V. alfalfae* (strain VaMs102) and *V. longisporum* (strain VL1) (Klosterman et al. 2011). In our analyses, we focused on the complete telomere-to-telomere genome assembly of *V. dahliae* strain JR2 (Faino et al. 2015). We first queried each of the 11,430 protein sequences of *V. dahliae* strain JR2 using BLAST against the aforementioned UniProtKB protein database. Subsequently, we utilized the Alien Index (AI) method (Gladyshev et al. 2008; Alexander et al. 2016) that generates AI scores for each *V. dahliae* gene based on the comparison of best BLAST hits with in-group and out-group species; in this case non-*Verticillium* fungal species and non-fungal species, respectively (Figure 2A). In this manner, we query for HGT candidates that were acquired only by *Verticillium* spp., or that were acquired by ancestral fungal species but kept only by *Verticillium* spp. during evolution. When the most significant BLAST hit of a gene within the out-group is more significant than the most significant hit within the in-group, the AI score is positive (i.e. AI>1) and this gene is considered a potential HGT candidate. This AI analysis yielded 42 HGT candidates with AI positive scores (AI>1) for *V. dahliae* JR2 (Figure 2B), which were further analyzed. To this end, we aligned protein sequences of all the homologs for each candidate (from in-, out-groups) and constructed phylogenetic trees that were automatically evaluated to remove phylogenetic trees with candidate genes that are directly adjacent to non-*Verticillium* fungal species (in-group species) rather than non-fungal species (out-group species). However, candidate genes that are closely related to the homologs of out-group species in a phylogenetic tree can also result from duplication followed by multiple gene losses in the in-group species. Typically, in such a phylogenetic tree, paralogs of the candidate gene cluster with genes of non-*Verticillium* fungal species (in-group species) and form separate branches from the branch of the candidate gene. Thus, we removed the candidates with phylogenetic trees that can be explained by gene losses in multiple species. In the end, we obtained two HGT candidates (HGT1 and HGT2) that were acquired only by *Verticillium* spp., or that were acquired by ancestral fungal species but kept only by *Verticillium* spp. during evolution. To check whether these two *V. dahliae* candidates are also present in other *Verticillium* spp., we queried them using BLAST against the proteomes of previously published *Verticillium* spp. (Shi-Kunne et al. 2018; Depotter et al. 2017). We found that candidate VDAG_JR2_Chr4g10560 (HGT1) is present in all *Verticillium* species and that VDAG_JR2_Chr8g03340 (HGT2) is only present in *V. alfalfae, V. nonalfalfae* and *V. longisporum* (Figure 3 & Figure 4).

**Figure 1.**
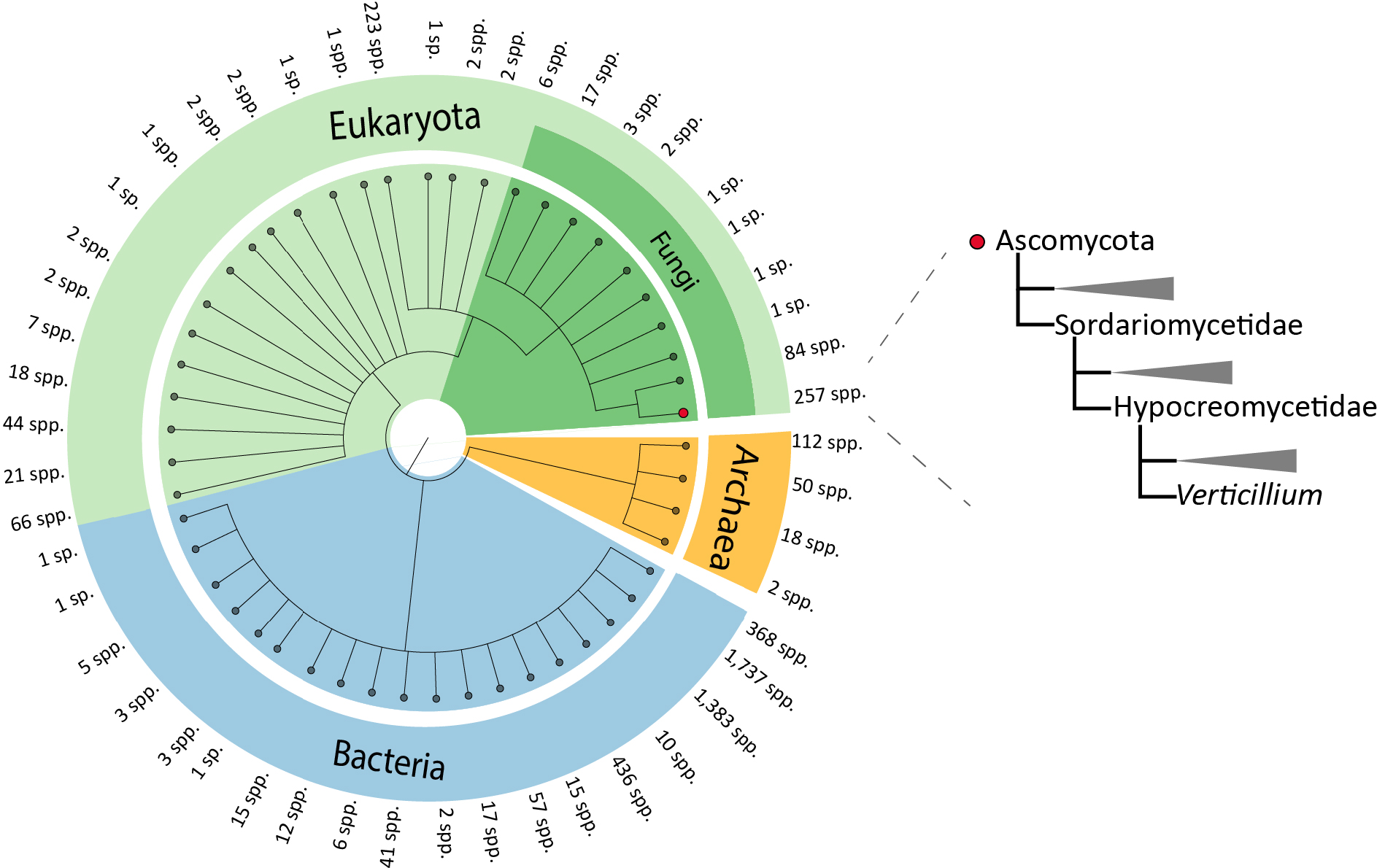
Phylogenetic tree of the species in the selected local data base. Different colors describe different taxonomical groups. The amount of species (spp.) on each branch is indicated. The phylogenetic position of *Verticillium* spp. is indicated on the right.

**Figure 2.**
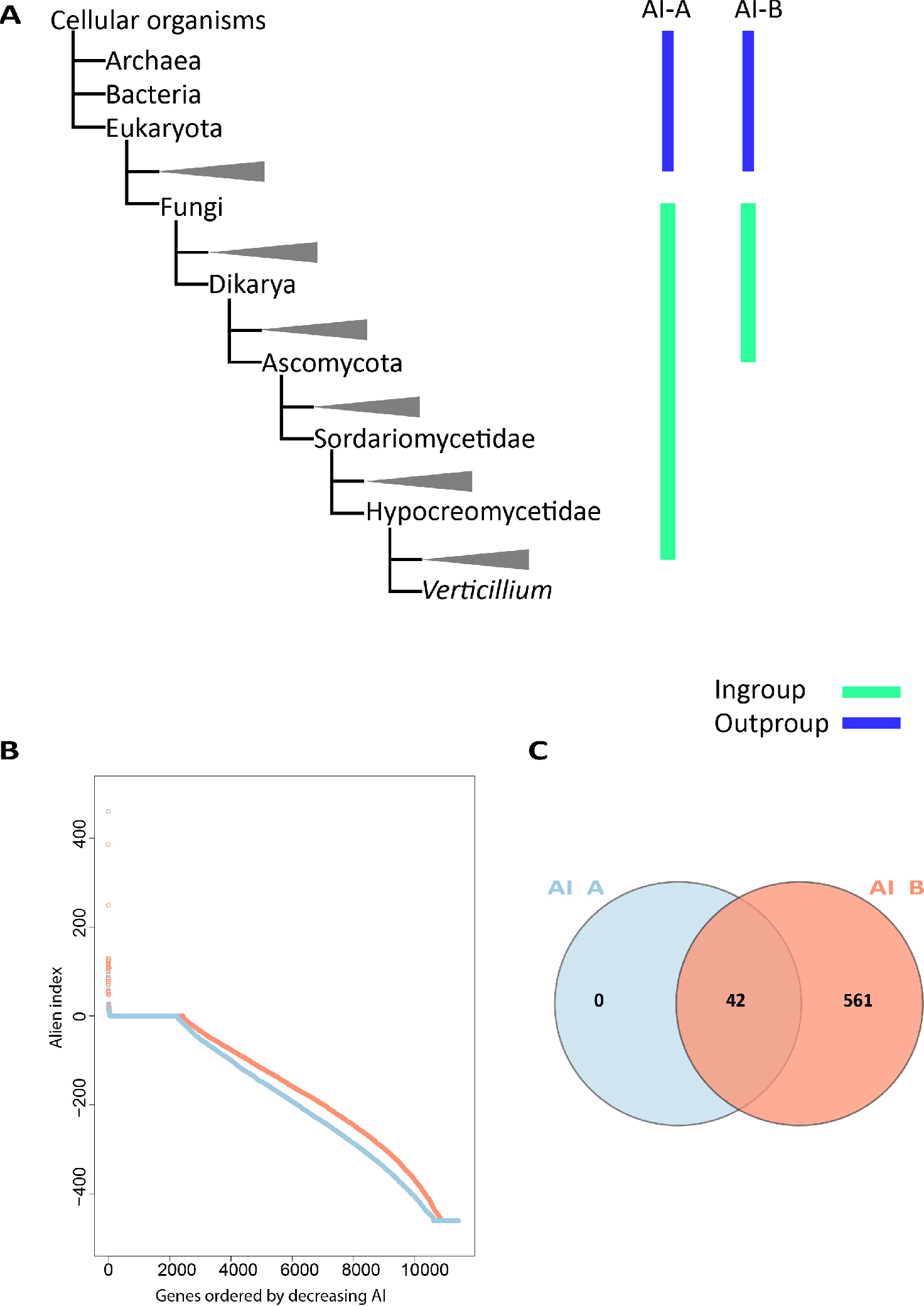
Alien Index based HGT detection method for detecting HGT candidates. (A) Simplified phylogenies of in- and out-group. In AI-A setting, in-group and out-group species are non-*Verticillium* fungal species and non-fungal species, respectively. In AI-B setting, in-group and out-group species are non-ascomycete fungal species and non-fungal species, respectively. (B) Distribution of AI sores. AI scores from two settings were calculated for every gene in the genome of *V. dahliae* strain JR2 and they were ordered by decreasing AI score. (C) Numbers of genes with AI positive scores from two different settings.

**Figure 3.**
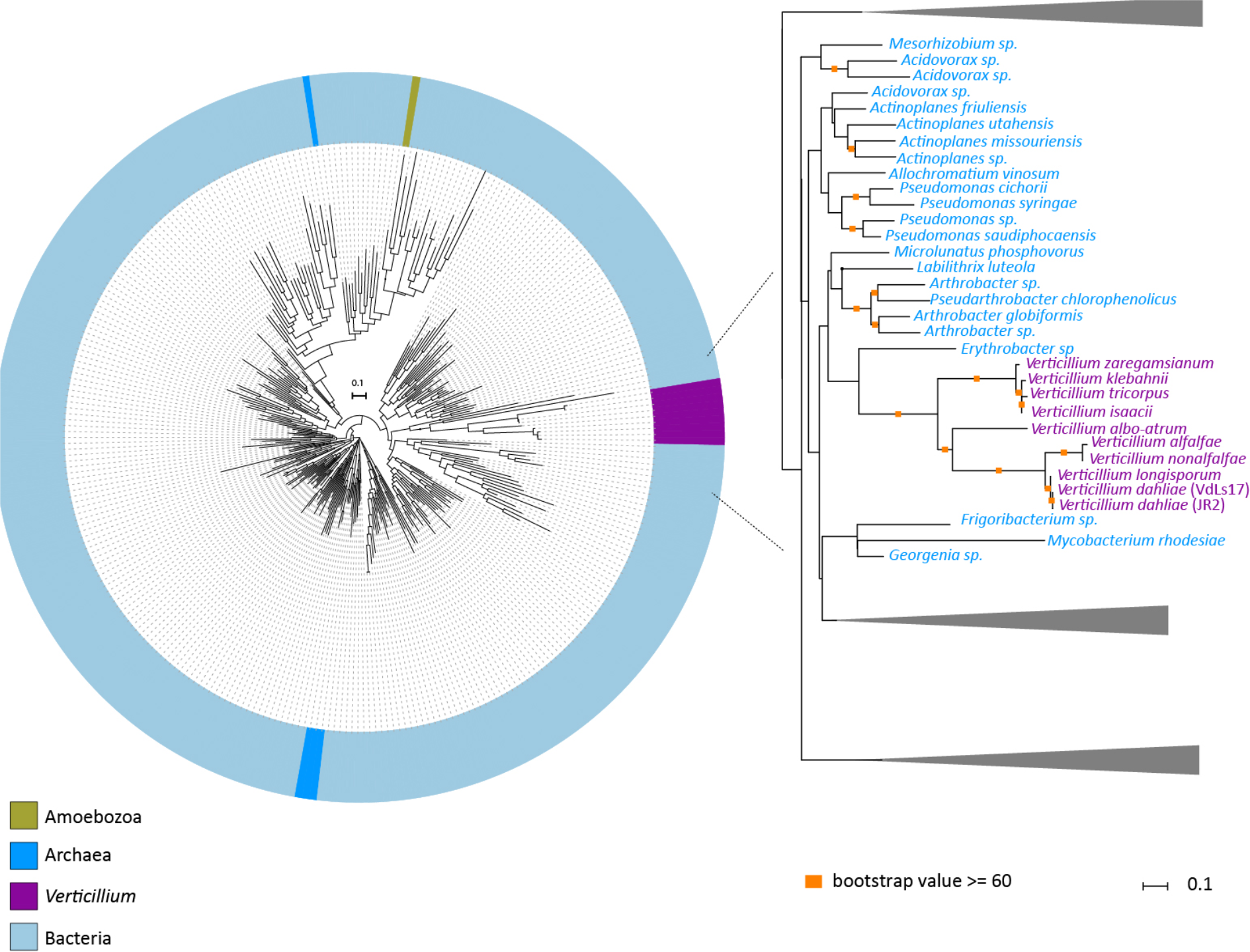
Evolutionary relationship of HGT-1 homologs. Protein sequences of HGT-1 orthologs were aligned and the resulting alignment was used to infer a maximum-likelihood phylogeny. The phylogeny suggests that *HGT-1* is transferred from a bacterial species. Different colors depict different groups or species. A more detailed part of the tree that contains *Verticillium* species is shown on the right. Orange squares indicate branches with bootstrap values ≥60.

**Figure 4.**
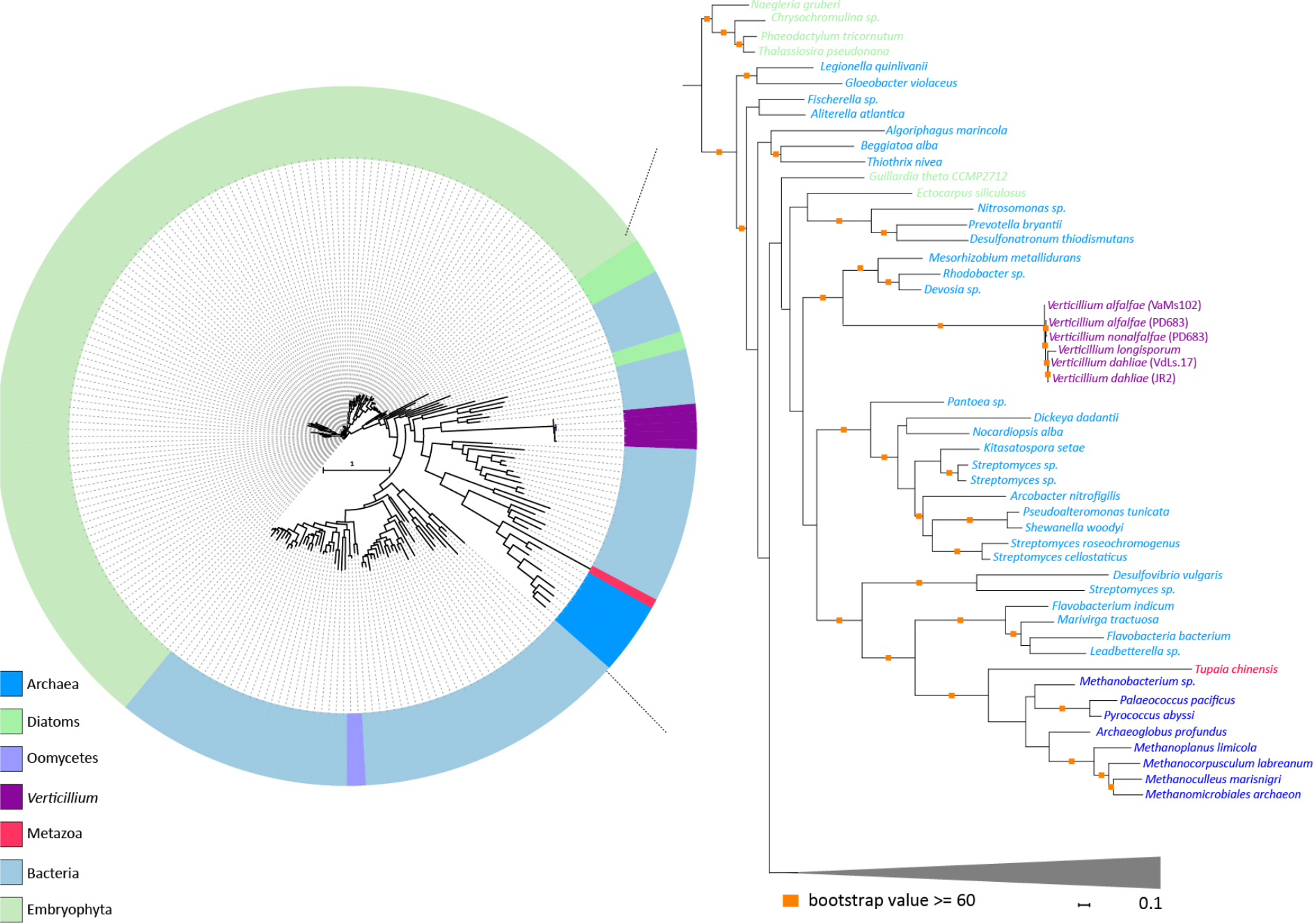
Evolutionary relationship of HGT-2 homologs. Protein sequences of HGT-2 orthologs were aligned and the resulting alignment was used to infer a maximum-likelihood phylogeny. Different colors depict different groups or species. The phylogeny suggests that *HGT-2* is transferred from a bacterial species. A more detailed part of the tree that contains *Verticillium* species is shown on the right. Orange squares indicate branches with bootstrap values ≥60.

In order to search for HGT candidates that are not only present in *Verticillium* spp., but also may be present in other ascomycete species, we again used the AI method, albeit with adjusted AI group settings. In this case, we set non-fungal species and non-ascomycete fungal species as out- and in- groups, respectively. AI scanning resulted in 603 genes with AI positive scores (AI>1), which were selected for further assessment through phylogenetic tree analysis (Table 1). After assessing phylogenetic trees, we identified 43 additional HGT candidates with non-fungal origins that are also present in other fungal ascomycete species. Among these candidates, two appeared as paralogs in the phylogenetic tree. The clusters of these two genes are close to each other in the tree, which indicates that the two paralogs are likely the result of a duplication and not of independent acquisitions. Therefore, we only considered 42 HGT candidates for further analyses. Of these, one is of plant origin and 41 are of bacterial origin (Table 1). The candidate of plant origin is the previously identified HGT gene, *Ave1* (Figure 5) (de Jonge et al. 2012). The previously reported glucosyltransferase gene (Klosterman et al. 2011) likewise was identified as HGT event in our study (Figure 6). Among these 42 candidates, 32 are present in all of the *Verticillium* species. Eight of the remaining ten are only found in the *Verticillium* species of the Flavnonexudans clade (Figure 7).

**Table 1.**
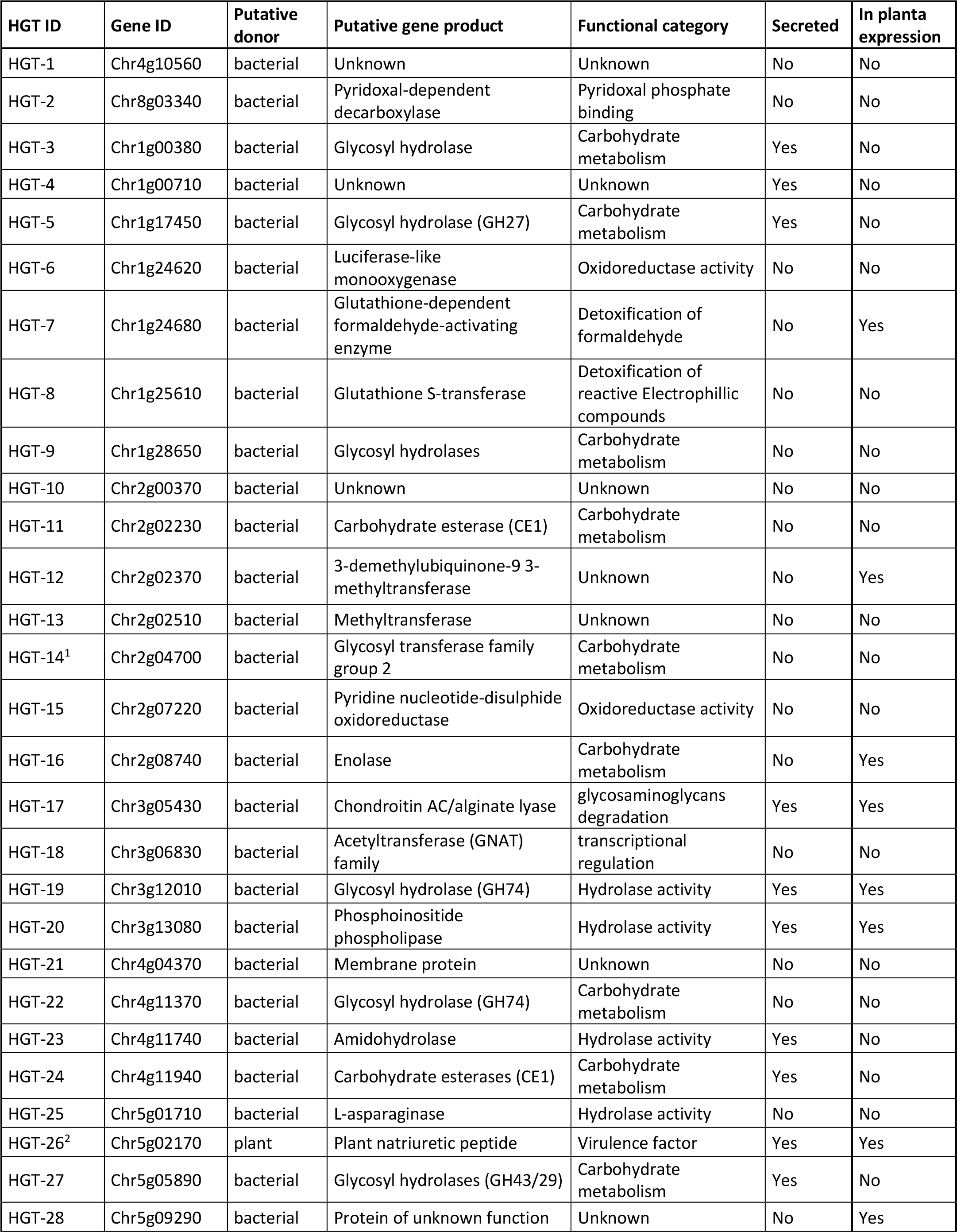
Information on HGT candidates.

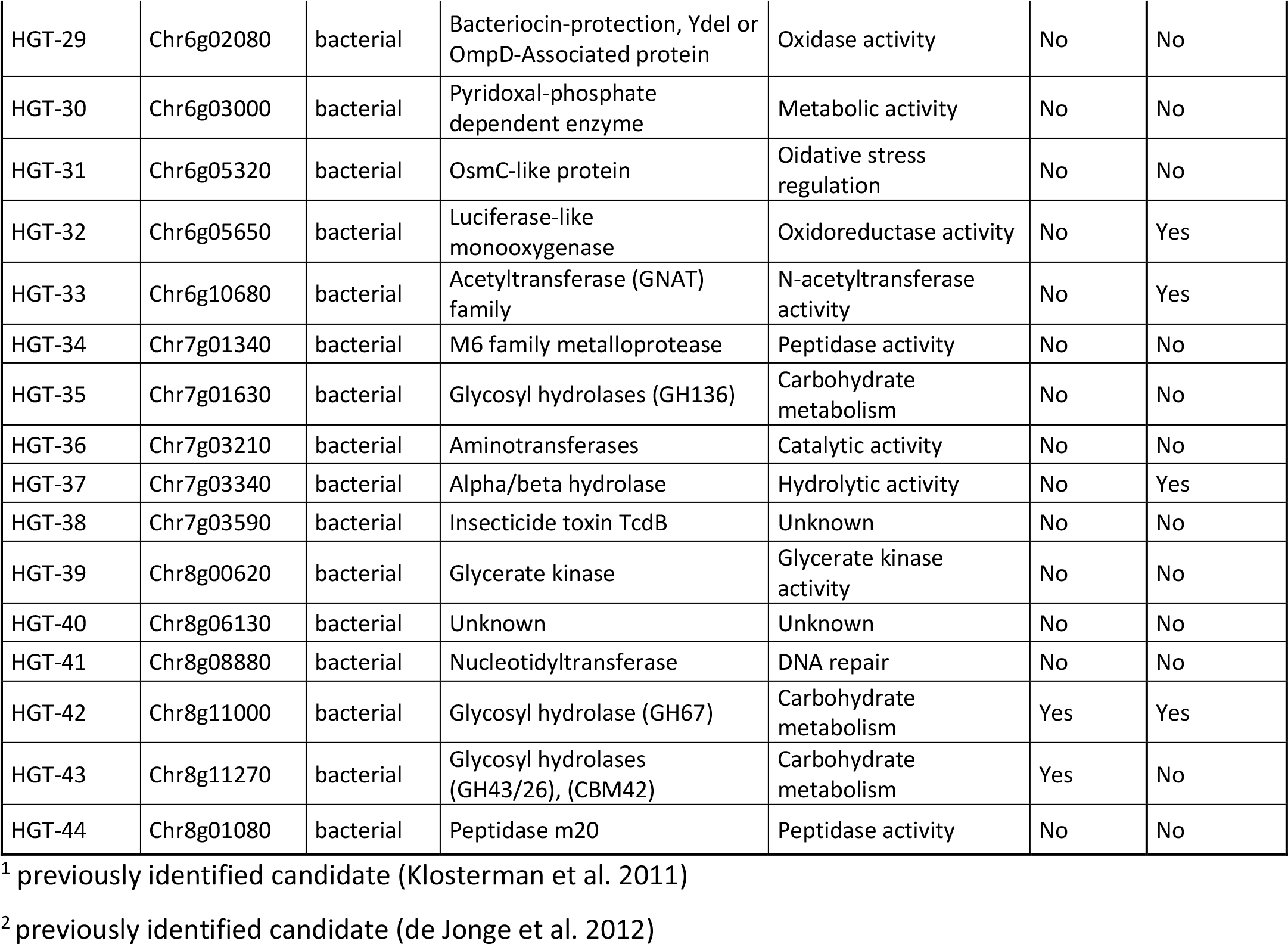

**Figure 5.**
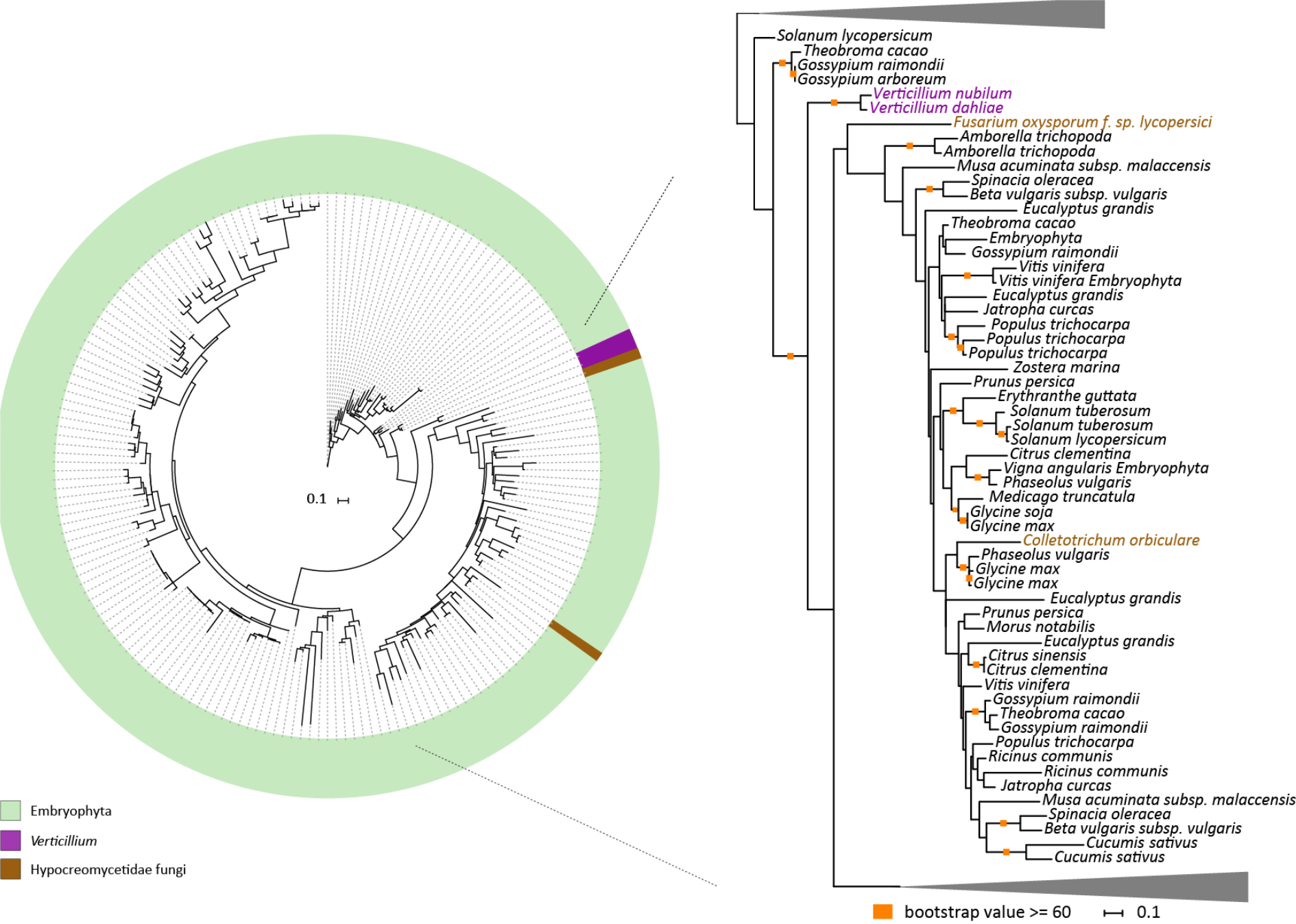
Evolutionary relationship of Ave1 homologs. Protein sequences of Ave1 orthologs were aligned and the resulting alignment was used to infer a maximum-likelihood phylogeny. Different colors depict different groups or species. A more detailed part of the tree that contains *Verticillium* species is shown on the right. Orange squares indicate branches with bootstrap values ≥60 or above.

**Figure 6.**
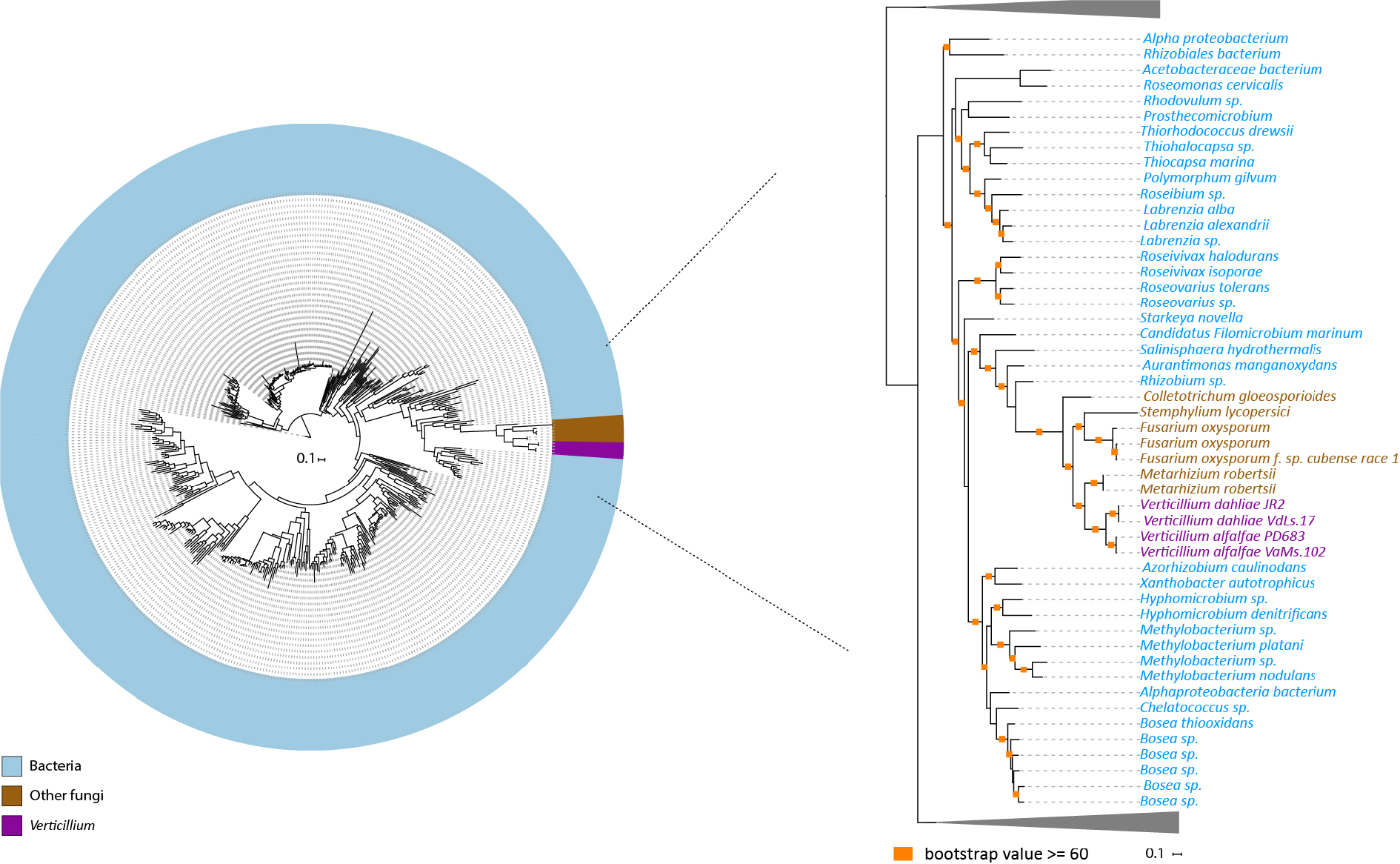
Evolutionary relationship of *V. dahliae* glucosyltransferase homologs. Orthologs of glucosyltransferase were aligned and the resulting alignment was used to infer a maximum-likelihood phylogeny. Different colors depict different groups or species. A more detailed part of the tree that contains *Verticillium* species is shown on the right. Orange squares indicate branches with bootstrap values ≥60.

**Figure 7.**
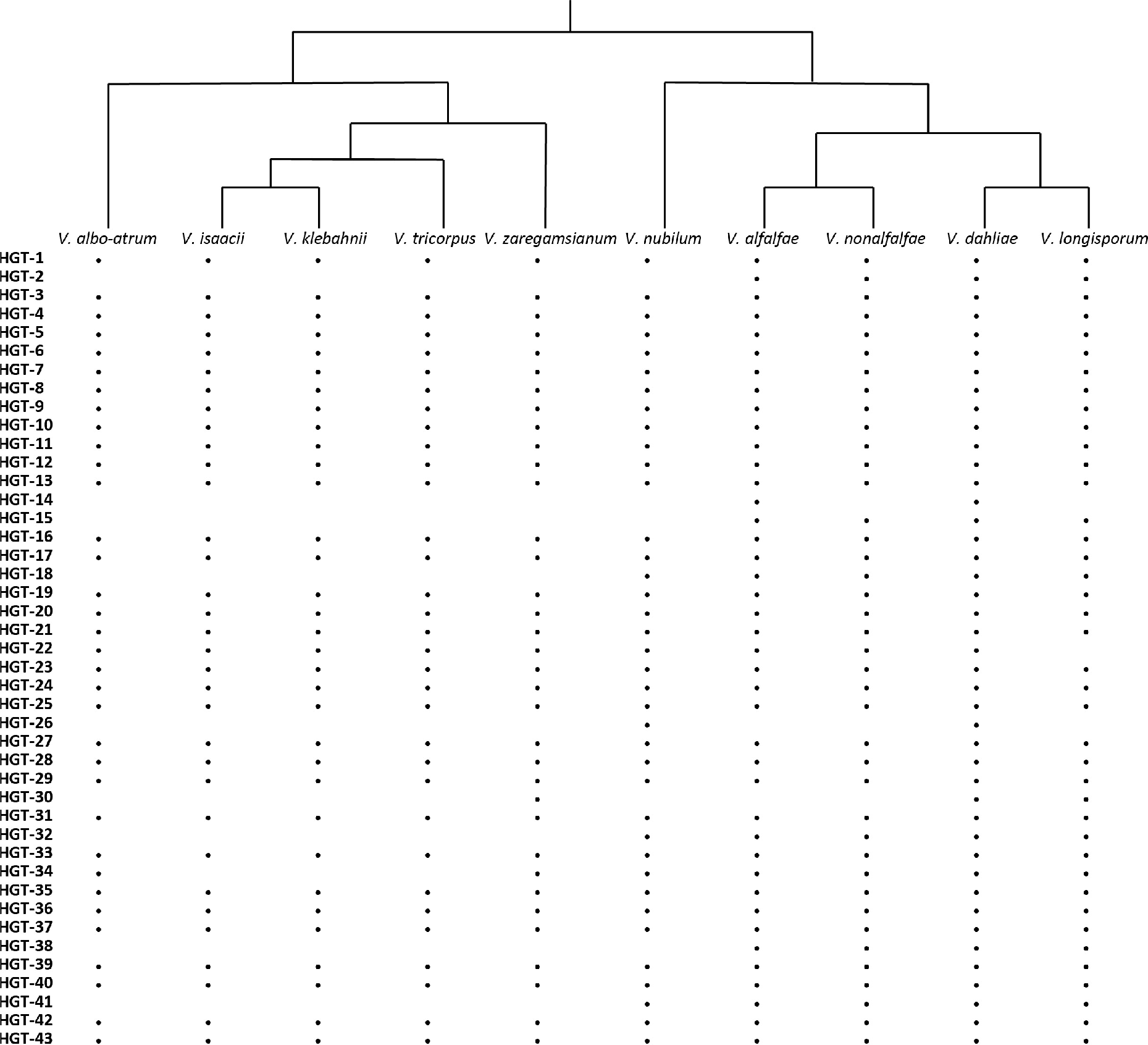
Presence and absence of all HGT candidates in all *Verticillium* species. Black dots indicate the presence of an HGT candidate in the corresponding *Verticillium* species.

### A high number of *V. dahliae* HGT candidates encodes secreted proteins

Filamentous plant pathogens secrete large numbers of proteins, including plant cell wall-degrading enzymes and effector proteins, to interact with host plants (Cook et al. 2015). We used a combination of SignalP, TMHMM and TargetP (Shi-Kunne et al., 2018) to identify HGT-derived genes that encode such secreted proteins. This analysis showed that 12 of the 44 candidates encode secreted proteins. Compared to the total number of the secreted proteins in the genome of *V. dahliae* strain JR2 (858 out of 11,430), HGT candidates are significantly enriched for genes that are predicted to encode secreted proteins (Fisher exact test, *p* < 0.0001). To know more about the putative functions of these proteins, we searched for conserved domains in each of these proteins using Interproscan (Jones et al. 2014). Among the 12 secreted proteins, five are glycoside hydrolases (GHs) and two are carbohydrate esterases (CEs), which are all carbohydrate-active enzymes (CAZymes). CAZymes are responsible for the synthesis and breakdown of glycoconjugates, oligo- and polysaccharides. CAZymes of plant associated fungi usually comprise a large number of plant cell-wall degrading enzymes (Zhao et al. 2013). The remaining five are Ave1, a chondroitin AC/alginate lyase, a phosphoinositide phospholipase, an amidohydrolase and a protein without functional annotation. Subsequently, we also searched for the functional annotation of the 30 non-secreted proteins. Besides the previously identified glucosyltransferase and seven proteins without functional annotation, we obtained diverse predicted functions including transcription regulation, DNA repair, hydrolase, and metabolic activities. Overall, we observed that the full set of HGT candidate proteins (secreted and non-secreted) is also enriched for GHs (10 out of 44) (Fisher exact test, *p* < 0.0001) when considering to the total amount of the GH genes in the genome of *V. dahliae* strain JR2 (265 out of 11,430).

To investigate the putative role of the HGT-derived genes in plant pathogen interactions, we compared expression patterns *in planta* and *in vitro*. It was previously shown that *Ave1* is expressed *in planta* during interaction of *V. dahliae* with *Nicotiana benthamiana* plants (de Jonge et al. 2012). We assessed transcription (RNA-seq) data of *V. dahliae* during colonization of *A. thaliana* and found that 12 HGT candidates showed *in planta* expression, one of which is *Ave1*. Four of these 12 were found to be induced when compared with the expression in *in vitro*-cultured mycelium (Figure 8). This suggests that besides *Ave1*, three additional HGT candidates might play a role during host colonization. Moreover, one of the three induced candidates encodes a secreted protein.

**Figure 8.**
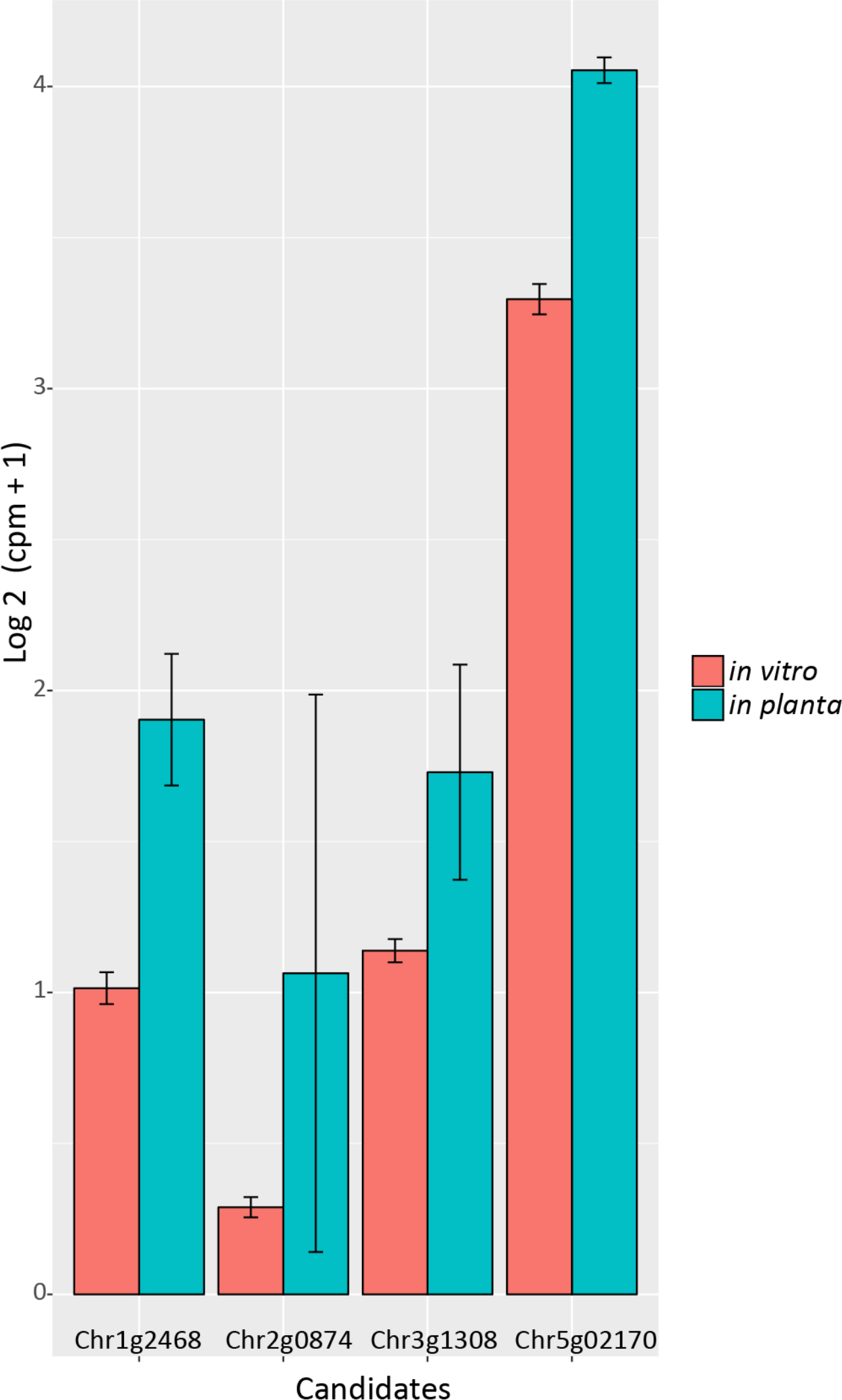
Pair-wise comparison of HGT candidates with differential expression *in vitro* and *in planta*. Gene expressions are depicted for *V. dahliae* strain JR2 cultured in liquid medium and upon *A. thaliana* colonization, respectively. Bars represent the mean gene expression with standard deviations. The significance of difference in gene expression was calculated using t-tests relative to a threshold (TREAT) of log2-fold-change ≥ 1 (McCarthy & Smyth 2009).

### *V. dahliae* HGT candidates mostly localize at repetitive genomic regions

In principle, foreign DNA can be incorporated anywhere in a recipient genome as long as it does not disrupt an essential genomic element (Husnik & Mccutcheon 2017). Several studies suggest that horizontally transferred sequences are often integrated into genomic regions that are enriched in transposable elements (TE) (Husnik & Mccutcheon 2017; Gladyshev et al. 2008; Mcnulty et al. 2010; Acuña et al. 2012; Pauchet & Heckel 2013). Previously, we observed that *Ave1* is located in a lineage-specific (LS) region of *V. dahliae* that is enriched in repeats (de Jonge et al., 2013; Faino et al., 2016). In order to assess whether the other HGT candidates are lineage-specific and associated with repeats as well, we plotted genomic repeat densities and the genomic location of all HGT candidates. Although only Ave1 is located at a genuine LS region (Depotter et al. 2018), most of the HGT candidates (34 out of 44) reside in proximity (within 1 kb) to a repetitive sequence (Figure 9). Subsequently, we compared repeat counts within 1 kb genomic windows around each of the 44 HGT candidates to the repeat counts in 1 kb genomic windows of 44 randomly picked genes (1000 permutations), which confirmed that the HGT candidates are significantly more often adjacent to repeats than expected by chance (*P* = 0.0331).

**Figure 9.**
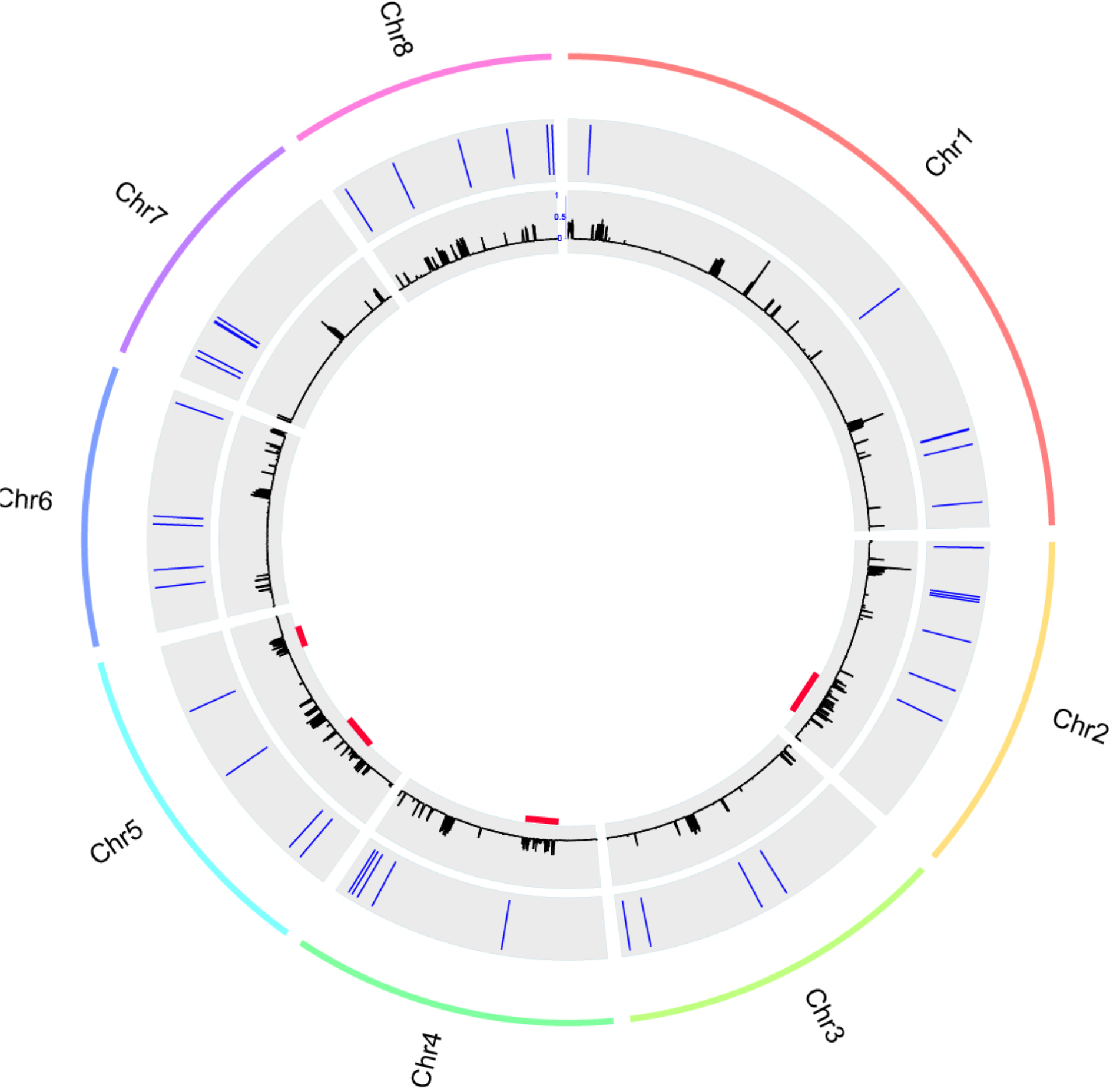
Genomic location of the HGT candidates. The outer lane represents the chromosomes of *V. dahliae* strain JR2. The middle lane shows the relative position of the HGT candidates in each chromosome. The inner lane shows the repeat density. The red lines indicate the locations of the lineage specific (LS) regions (Depotter et al. 2018).

## DISCUSSION

In this study, we systematically searched for evidence of inter-kingdom HGT events in the genome of *V. dahliae* using an Alien Index (AI) based method. Subsequently, we verified HGT candidates using a phylogenetic approach. In general, there are two classical ways to identify HGT candidates: intrinsic and extrinsic methods. Intrinsic methods focus on patterns in the primary structure of genes and genome sequences and aim to find genes or genomic regions with composition patterns that differ significantly from the rest of the genome. For example, deviation in GC content and codon usage can be considered a sign for horizontal acquisition (Lawrence & Ochman 1997). In contrast, extrinsic methods focus on comparing similarity metrics between closely related and distant taxa. For example, when a gene from a species of interest (recipient) shows higher similarity (lower E-value, higher bit-score in BLAST) to sequences from distantly related (donor) species than to genes of close relatives, this gene may be horizontally acquired. Another frequently applied extrinsic approach is to assess the phylogenetic distribution of genes across a panel of diverse organisms. These phylogeny-based methods are generally thought to be more accurate (Poptsova & Gogarten 2007; Poptsova 2009). In principle, an HGT event will create a discrepancy between the gene and species trees. However, phylogeny-based methods are much more computationally intensive, because they require ortholog predictions, sequence alignments and phylogeny reconstructions (Dupont & Cox 2017). Thus, we chose to utilize an AI based method to pre-select candidates that can be further analyzed through phylogenetic analyses.

Although the AI method provided a rapid identification of HGT candidates, some limitations of the method should be kept in mind. The AI method relies on the quality of the reference database, the accuracy of the taxonomic assignment and the diversity of taxa covered by the reference database. The two HGT candidates (HGT1 and HGT2) that appear to be *Verticillium*-specific could also be explained by the absence of the ortholog-containing species. In other words, the more closely related species are included in the database, the less *Verticillium*-specific HGT candidates might be identified. Furthermore, AI based methods rely on pre-defining in- and out- groups and can only identify HGT candidates that are absent in the in-group species. In our case, the method is unable to identify candidates that were acquired before the formation of ascomycetes, and candidates that were also acquired by fungal non-ascomycete species independently at more recent events from the same donors.

Nevertheless, the 42 HGT candidates that were also found in other ascomycete species were most likely acquired during ancient transfer events. The extent of gene transfer from prokaryotes to fungi is highly different across diverse ascomycete fungal lineages. A whole-genome study of HGT in *Aspergillus fumigatus* revealed that 40% of 189 transferred regions (containing 214 genes) were of bacterial origin (Mallet et al. 2010). In contrast, a study with a similar objective found only 11 genes with a convincing signature of transfer from bacterial origin in the genomes of three *Colletotrichum* species (Jaramillo et al. 2015). Three of the 11 *Colletotrichum* HGT candidates were also reported to be found in *V. dahliae* (strain VdLs17) and *V. alfalfae* (strain VaMs102) (Klosterman et al. 2011). In our study, we also found these three candidates (HGT-14, HGT-15 and HGT-44) in *V. dahliae* strain JR2.

We found that HGT candidates are enriched for genes encoding secreted proteins, many of which belong to a family of carbohydrate-active enzymes (CAZymes) or glycoside hydrolases (GHs). GHs play a fundamental role in the decomposition of plant biomass (Murphy et al. 2011), which suggests that bacterial genes might contribute to the plant-associated life styles of *Verticillium* species as well as of many other ascomycetes. Although these *V. dahliae* GH genes are not induced during infection of *A. thaliana* when compared with *in vitro* expression data, it needs to be noted that *V. dahliae* is able to cause disease in hundreds of plant species (Inderbitzin & Subbarao 2014; Fradin & Thomma 2006), and it might be possible that the GH genes are induced during interaction with other host plants. Alternatively, these proteins might play roles during life stages outside the host. Overall, when assessing transcription (RNA-seq) data of *V. dahliae*-infected *A. thaliana*, we identified four HGT candidates (including *Ave1*) that were induced when compared with expression in *in vitro*-cultured mycelium. One of the induced candidates encodes a secreted protein that is predicted to be a phosphoinositide phospholipase, a key metabolic enzyme that is needed by all living organisms to hydrolyze phospholipids into fatty acids and lipophilic substances (Ghannoum 2000). Phospholipases are also signaling molecules that elicit stress tolerance and host immune responses in fungi (Ghannoum 2000; Köhler et al. 2006; Soragni et al. 2001). Phospholipases have been studied in some plant pathogenic fungi, such as *Fusarium graminearum* (Zhu et al. 2016) and *Magnaporthe oryzae* (Zhang et al. 2011; Ramanujam & Naqvi 2010; Yin et al. 2016). The phospholipase (FgPLC1) is considered to be involved in regulation of development, stress responses and pathogenicity of *F. graminearum* (Zhu et al. 2016). Collectively, this may suggest that the *V. dahliae* phosphoinositide phospholipase might play an import role during interaction with host plants.

## CONCLUSIONS

In this study, we applied a conservative approach combining an Alien Index (AI) method and phylogenetic analysis to analyze inter-kingdom HGT to *Verticillium dahliae*. Besides the previously identified effector gene *Ave1* and a glucosyltransferase-encoding gene, we revealed 42 additional HGT candidates, all of which are of bacterial origin, indicating a high number of inter-kingdom gene acquisitions.

## MATERIALS AND METHODS

### Construction of local protein database and protein BLAST

The protein database used for BLAST analyses was generated using the reference proteomes from UniProtKB excluding viruses (downloaded at 07-07-2016). Entries from UniProt were renamed to include their UniProt ID and NCBI taxonomy ID. Sequences shorter than ten amino acids were removed from the database. All predicted proteins sequences from each of the *Verticillium* species were queried to search homologs in the local protein database using BLASTp (with default settings). We searched for the presence of the HGT candidates in one strain per species that were studied previously (Depotter et al., 2017; Shi-Kunne et al., 2018). Protein sequences of each HGT candidate were queried against the proteome of each strain using BLASTp (with default settings).

### Identification of HGT candidates

Alien Index (AI) scores were calculated using a custom-made python script which applied the following formula: *AI* = (ln (*bbhG* + 1 × 10^−200^) − ln (*bbhO* + 1 × 10^−200^)), where bbhG and bbhO are the E-values (<10^−3^) of the best BLAST hit from the in-group and out-group, respectively. Best BLAST hits were determined by the highest bit score. Genes with AI score >1 were classified as positive AI genes. For each AI positive gene, homologs were selected based on the BLAST criteria that at least 70% of query length is covered by at least 60% of hit sequence using a custom-made Python script. These homologs were then aligned using MAFFT with default setting (Katoh & Standley 2013). Alignments were subsequently curated using Gblocks (Castresana 2000) with non-stringent parameters. Phylogenetic trees of the curated alignments were constructed using FastTree (Price et al. 2010) with gamma likelihood and wag amino acid substitution matrix. Phylogenetic tress were automatically evaluated using a custom made Python script that removes phylogenetic trees with candidate genes that are directly adjacent to non-*Verticillium* fungal species (in-group species) rather than non-fungal species (out-group species). Subsequently, new phylogenetic trees (for manual inspection) were reconstructed using RAxML (Stamatakis 2014) with automatic substitution model determination for branching values.

### Gene expression analysis

To obtain RNA-seq data for *V. dahliae* grown in culture medium, isolates JR2 was grown for three days in potato dextrose broth (PDB) with three biological replicates. To obtain RNA-seq data from *V. dahliae* grown *in planta*, three-week-old *A. thaliana* (Col-0) plants were inoculated with isolate JR2. After root inoculation, plants were grown in individual pots in a greenhouse under a cycle of 16 h of light and 8 h of darkness, with temperatures maintained between 20 and 22°C during the day and a minimum of 15°C overnight. Three pooled samples (10 plants per sample) of complete flowering stems were used for total RNA extraction. Total RNA was extracted based on TRIzol RNA extraction (Simms et al. 1993). cDNA synthesis, library preparation (TruSeq RNA-Seq short insert library), and Illumina sequencing (single-end 50 bp) was performed at the Beijing Genome Institute (BGI, Hong Kong, China). In total, ∼2 Gb and ∼1.5 Gb of filtered reads were obtained for the *V. dahliae* samples grown in culture medium and *in planta*, respectively. RNAseq data were submitted to the SRA database under the accession number: SRP149060.

The RNA sequencing reads were mapped to their previously assembled genomes using the Rsubread package in R (Liao et al. 2013). The comparative transcriptomic analysis was performed with the package edgeR in R (v3.4.3) (Robinson et al. 2010; McCarthy et al. 2012). Genes are considered differently expressed when P-value < 0.05 with a log2-fold-change ≥ 1. P-values were corrected for multiple comparisons according to Benjamini and Hochberg (Benjamini & Hochberg 1995).

## ACKNOWLEDGEMENTS

Work in the laboratories of B.P.H.J.T. and M.F.S is supported by the Research Council Earth and Life Sciences (ALW) of the Netherlands Organization of Scientific Research (NWO).

